# Protal: Ultra-fast metagenomic profiling and strain-resolved analysis

**DOI:** 10.64898/2026.08.03.742433

**Authors:** Joachim Fritscher, Anthony Duncan, Falk Hildebrand

## Abstract

Large-scale metagenomic studies increasingly require taxonomic profiles that are sensitive, precise, strain-resolved and computationally tractable. Existing profilers typically trade taxonomic breadth, sensitivity, precision and speed against one another, limiting their utility for high-resolution microbiome analyses. Here we present *protal* — profiling through alignment — an ultra-fast alignment-based method for species-and strain-resolved profiling of metagenomes.

Protal combines a newly developed alignment algorithm, machine-learning-based classification and conserved bacterial marker genes to profile species represented in the standardized and regularly updated GTDB taxonomy. Protal reliably profiles all 143,614 bacterial and archaeal species in GTDB r226 and achieved higher precision on CAMI2 benchmarks than all tested contemporary profilers, including MetaPhlAn 4, mOTUs4, sylph and Kraken2+Bracken (mean species-level precision 98.3% versus 97.5% for the next-best profiler, sylph). In custom benchmarks, protal showed particularly strong gains for rare species represented by a single reference genome and for highly complex communities containing 10,000 species (F1-score 9% and 14% higher than the second best profiler, respectively). At the strain level, protal reconstructs intraspecific phylogenetic relationships among detected bacteria with similar accuracy as StrainPhlAn 4; because protal produces precise alignments, the phylogenies can be *de novo* produced without reliance on reference strain collections. Unlike dedicated strain-profiling workflows, however, protal performs strain analysis concurrently with species-level profiling, making it up to 40-fold faster without requiring additional steps. Together, these features make strain-resolved profiling of thousands of metagenomes feasible on commodity hardware. The software, databases and tutorials are available at https://github.com/4less/protal and http://protal.earlham.ac.uk.

## Introduction

Recent advances in metagenome assembly[1, 2] and genome binning[3, 4], together with the rapid accumulation of publicly available sequencing data, have dramatically expanded the diversity represented in reference genome databases such as the Genome Taxonomy Database (GTDB) [5].

Assembly-based approaches provide high-resolution characterization of microbial communities but remain computationally demanding at large scale and can fail to recover low-abundance taxa. Instead, reference-based taxonomic profiling offers a complementary approach that capitalizes on the expanded reference databases derived from both isolate genomes and MAGs. It is in general computationally more efficient and has an increased sensitivity for low-abundance taxa. These strengths make reference-based profiling particularly well-suited for analyzing well-described microbiomes such as the human gut, complex microbial communities with a large number of low-abundant taxa, and large-scale datasets.

Current reference-based profilers generally rely either on phylogenetically informative marker genes (e.g. MetaPhlAn 4[6] and mOTUs4[7]) or on genome-specific k-mers whose accumulated counts are taken as evidence for a species’ presence (e.g. Kraken2[8] and Sylph[9]).

Although strain-level variation is increasingly recognized as biologically important[10], most taxonomic profilers remain optimized for species-level inference. Strain-level analyses typically require separate downstream workflows based on variant calling or marker reconstruction, such as StrainPhlAn 4[11] for MetaPhlAn 4, and metaSNV [12] for mOTUs4, increasing computational complexity and limiting scalability.

To address the growing need for fast, highly resolved and precise metagenomic profilers, we introduce protal (profiling through alignment). Protal introduces a phylogenetically informed alignment framework that exploits unique GTDB marker genes to jointly infer taxonomic composition and strain relationships. Its performance is centered around a newly developed alignment algorithm, that is both more precise and faster than contemporary aligners. The key features of protal are (1) full compliance with the GTDB database and its updates, (2) robust representation of species with as little as one reference genome available, (3) evolutionary accurate strain-level phylogenies, and (4) one to two orders of magnitude faster profiling than comparable tools.

## Methods

### Protal database construction and alignment algorithm

Protal builds taxonomic reference databases from GTDB species-representative marker-genomes. Unless stated otherwise, our analyses used GTDB r226 [5] containing 143,614 species. Further, the GTDB universal marker-gene set was used as provided in the database [13, 14], comprising up to 120 bacterial and 53 archaeal universal single-copy marker genes per species. Further, we generated a protal-mini database containing 20 marker genes from both bacteria and archaea.

Protal uses a custom alignment algorithm based on two integrated k-mer representations, core-mer and flex-mer. A core-mer is the central 15-mer of a 31-mer and is used for exact lookup. A flex-mer is the 16-nt sequence formed by the two 8-nt flanks surrounding the core-mer and is used to rank candidate matches with identical core-mers. Core-mers and flex-mers are stored in a custom hash-based data structure, the flex-map, which records marker-gene location, taxon identity and k-mer uniqueness information. Protal further annotates short-unique k-mers, long-unique k-mers and long-super-unique k-mers, defined by increasing degrees of species specificity across the GTDB reference collection. These uniqueness features are not used to alter the alignment itself, but are recorded for downstream taxon classification.

For each read, protal extracts open syncmers from 15-mer core-mers [15] and subsequently queries them against the flex-map and retrieves candidate marker-gene locations. Candidate seeds are grouped into anchors by taxon and marker gene, ranked by the number of exactly matching positions and extended into gapped alignments using WFA2 [16, 17] with mismatch, gap-opening and gap-extension penalties of 4, 6 and 2, respectively. For paired-end reads, protal jointly evaluates compatible mate alignments and reports the best-scoring alignment pair. Protal outputs alignments in SAM format together with mapping quality, alignment identity and k-mer uniqueness features.

### Taxon detection and abundance estimation

Species detection is performed using a random-forest classifier[18] (512 trees, maximum of 256 leaf nodes/tree, class weight ‘balanced‘) trained on synthetic metagenomes with known source composition. For each species with at least one assigned read, protal records per-species alignment-derived features, including the number and distribution of detected marker genes, long-unique and long-super-unique k-mer counts, expected gene presence, mean mapping quality, mean alignment identity and coverage variation across marker genes. The number of variables considered at each split was tuned by 5-fold cross-validation. Training data were generated using simulate metagenomes from annotated GTDB r226 genomes: six synthetic datasets totaling 581 samples spanning 1–500 species, varying species richness, sequencing depth, abundance distribution and strain composition (full parameters in Supplementary Methods).

For each detected species, abundance was estimated from marker-gene read depth. Per-gene read depth was calculated as the total aligned read length divided by marker-gene length, and the species-level abundance estimate was taken as the median read depth across marker genes assigned to that species.

### Reference-guided strain-resolved alignments and phylogenies

For species detected across multiple samples, protal reconstructs one reference-guided marker-gene multiple sequence alignment (MSA) per species. Alignments with MAPQ *≤* 4 or soft-clip-corrected aligned length below 50 bases were discarded. At each marker-gene position, protal required a minimum vertical coverage of 2 to call a consensus base. For multi-allelic positions, the allele with the highest summed base quality was chosen if it had coverage *≥* 2 and either a summed Phred base-quality score *≥* 90 or a mean base quality *≥* 15. Positions without sufficient evidence were encoded as ambiguous or missing bases according to filtering rules described in Supplementary Methods.

Reference-guided MSAs were constructed by placing per-sample consensus bases relative to the marker-gene reference. Insertions were handled in two passes: first, protal identified the maximum insertion length at each reference position across samples; second, non-inserting samples were padded with gaps to preserve column alignment. This reference-guided strategy is conceptually similar to approaches used in ViralMSA [19] and VIRULIGN [20], scaling linearly with sample number.

Per-species MSAs were filtered using qcmsa.py (default preset). Sample–gene cells were retained if horizontal coverage was *≥* 0.3 and mean depth was *≥* 1.0*×*, and a gene was retained only if more than three samples passed this filter. Potential mixed-strain or contaminated samples and unreliable genes were then removed using an iterative Tukey interquartile-range outlier rule applied to per-gene and per-sample multi-allelicity rates. By default, qcmsa.py retains constant (invariant) positions, as they inform branch-length estimation; low-parsimony sites (positions at which fewer than two samples differ from the majority base) and sequences retaining fewer than 1000 aligned bases were removed before tree inference (Supplementary Methods). Phylogenetic trees were inferred from filtered MSAs using IQ-TREE (v3.1.1)[21] with the GTR model and fast search mode.

### Read-alignment benchmark

To benchmark alignment accuracy, reads were simulated from all GTDB r214 genomes, including non-representative genomes (*n* = 394,932), using ART Illumina 2.5.8 [22]. Simulations used 150-bp paired-end reads, HiSeq 2500 error profiles, mean insert size 200 bp and insert-size standard deviation 10 bp. Reads were aligned against GTDB r214 species-representative marker genes using protal, Bowtie2 2.5.3 [23], BWA-MEM2 2.2.1 [24], minimap2 2.28-r1209 [25] and Strobealign 0.13.0 [26]. Alignments were counted as true positives when the read mapped to a marker gene from the species cluster of origin and as false positives otherwise. True-positive and false-positive rates were calculated across mapping-quality thresholds.

Alignment runtime and memory benchmarks were run on an exclusively reserved compute node with an AMD EPYC 7713P CPU and 512 GB RAM, by running software 5 times and taking the result from the final run.

### Species-level profiling benchmarks

Species-level profiling was evaluated using CAMI synthetic metagenomes [27, 28], a rare-species benchmark and a high-complexity simulation. CAMI datasets included the Toy Human Microbiome Project datasets representing five body sites, the Toy Mouse Gut dataset and the Marine dataset. CAMI gold standards were converted to GTDB r226 by assigning source genomes with GTDB-Tk [29] and transferring the corresponding abundance profiles.

The rare-species benchmark (R.Sp.) was designed to evaluate detection of species represented by only one genome in GTDB r214. We selected 958 isolate genomes from 364 such species from the HumGut database [30]; these genomes were subsequently re-annotated with GTDB-Tk against GTDB r226 for evaluation. We simulated 20 metagenomes, each containing 200 species represented by one randomly selected genome. Reads were simulated using ART Illumina with 150-bp paired-end reads, mean insert size 200 bp and insert-size standard deviation 10 bp.

The high-complexity metagenome benchmark contained 10,000 species per simulated metagenomic sample and was designed to evaluate profiler robustness under extreme species richness. 10 samples were simulated with species randomly selected from GTDB r226.

Protal v0.6.0 (run in both its full and reduced “protal-mini” database configurations) was compared with MetaPhlAn 4 v4.2.4 [6], mOTUs v4.0.4 [7], Kraken2 v2.17.1 with Bracken v3.1 [8, 31], and sylph v0.9.0 [9]. MetaPhlAn 4 was run with database mpa vJan21 CHOCOPhlAnSGB 202103. Kraken2 was run against a custom database built from all GTDB r226 species-representative genomes, with Bracken abundance estimation using threshold-t 450. Sylph was run with GTDB r226 database built using either *c* = 200 or *c* = 1000 as sketching parameter.

Predictions that a profiler explicitly declined to name at the rank being evaluated were excluded from scoring at that rank, rather than counted as false positives. This applies to MetaPhlAn 4’s UNCLASSIFIED fraction and to mOTUs4’s unnamed marker-gene clusters (reported as s__Unknown <genus> mOTUv4.0_*), which are resolved to genus but carry no species name and therefore cannot match a gold-standard species. Such predictions still contribute at ranks where they are named, so genus-and higher-level results are unaffected.

Profiler outputs were converted to GTDB taxonomy, where required. MetaPhlAn 4 was mapped using sgb to gtdb profile.py; mOTUs output was mapped using the corresponding mOTUs-to-GTDB mapping file; and Kraken2+Bracken output was native to GTDB r226 and required no mapping.

### Strain-level phylogeny benchmark

We generated the STRAIN46 dataset to evaluate strain-resolved phylogeny reconstruction. Source genomes were selected from HumGut [30]. Isolate genomes from 50 species with 15–60 intraspecific genomes each were re-annotated with GTDB-tk (GTDB r226). After removal of species with fewer than 15 genomes after re-annotation, the final dataset comprised 1,346 genomes across 46 species.

We simulated 200 metagenomes, each containing one strain (genome) from each of the 46 species. Strains were selected randomly, and vertical abundance was drawn from a negative-binomial distribution scaled between 1 and 50. Reads were simulated using ART Illumina with 150-bp paired-end reads, HiSeq 2500 error profiles, mean insert size 200 bp and insert-size standard deviation 10 bp. Gold-standard core-genome alignments were generated using Roary[32] v3.13.0 with --mafft [33] on Prokka[34] gene predictions; gold-standard phylogenies were inferred using IQ-TREE [21] with the GTR model.

Protal strain-level output was compared with MetaPhlAn 4+StrainPhlAn 4 (v4.2.4) [6]. StrainPhlAn 4 marker regions were extracted using sample2markers.py and extract markers.py, and per-species MSAs were generated using StrainPhlAn 4.

### Benchmark metrics

Species detection was evaluated using sensitivity, precision and F1-score. Abundance estimation was evaluated using Bray–Curtis dissimilarity, L2 error and Pearson correlation between predicted and expected abundance vectors. Metrics were calculated using custom scripts available at https://github.com/4less/benchpro_rust.

Strain-level performance was evaluated using MSA error, strain monophyly and tree similarity to gold-standard phylogenies. MSA error was calculated by comparing reconstructed marker-gene consensus sequences with the corresponding source genome sequence. For monophyly, samples simulated from the same source genome were expected to form a monophyletic group. We calculated monophyly as the number of tips from the source genome divided by the total number of tips under their lowest common ancestor. Tree similarity was calculated after restricting inferred and gold-standard trees to shared tips, requiring at least four shared tips. We quantified tree disagreement using branch-score distance [35] and path difference [36].

## Results

### Protal implementation

Protal uses universal single-copy marker genes as references, which support accurate species detection and abundance estimation while also providing conserved evolutionary signal for strain-resolved analyses. Metagenomic reads are aligned against marker genes to derive both taxonomic profiles and strain-resolved phylogenies. For example, using GTDB r226 [5], the protal reference database represents 143,614 species (Fig. 1a,b).

**Fig. 1.**
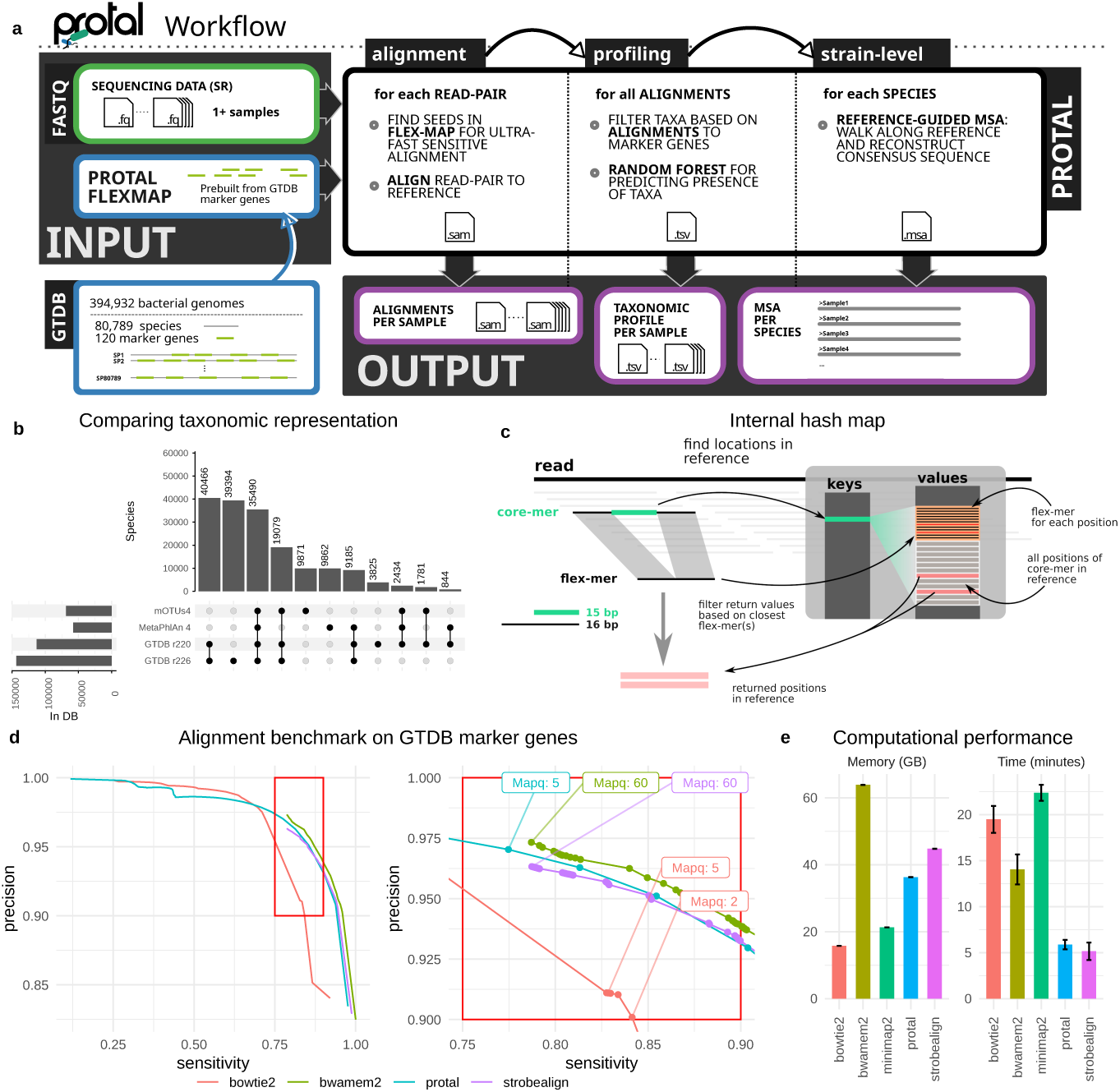
Overview of protal workflow, database coverage, alignment strategy and alignment performance. **a** Protal takes single-or paired-end short-read metagenomic data from one or more samples as input. Its main outputs are read-to-marker-gene alignments in SAM format, taxonomic profiles for each metagenome, and, for each detected species, marker-gene multiple sequence alignments reconstructed across samples. **b** Species-level coverage of protal databases built from GTDB releases r220 and r226, compared with the taxonomic coverage of mOTUs4 and MetaPhlAn 4. **c** The flex-map data structure stores core-mers representing exact seed matches, flex-mers representing tolerated inexact matches, and taxonomic identifiers. This structure enables sensitive alignment while reducing memory use and maintaining high mapping speed (see also Suppl. Fig. S5). **d** Alignment performance of protal, Bowtie2, BWA-MEM2, minimap2 and Strobealign. True-positive rate and false-positive rate were calculated across mapping-quality thresholds. All tools were run against the same reference database of representative GTDB marker genes, and reads were simulated from entire GTDB r214 genomes. **e** Runtime and peak memory use for aligning 2 × 7.9 GB of uncompressed paired-end reads to the representative GTDB marker-gene reference database used in **d**.

During read alignment, protal records species-specific unique k-mers and additional alignment-derived features. These features are subsequently evaluated by a random forest classifier (Fig. 2e) to determine which species are present and to estimate their relative abundance.

**Fig. 2.**
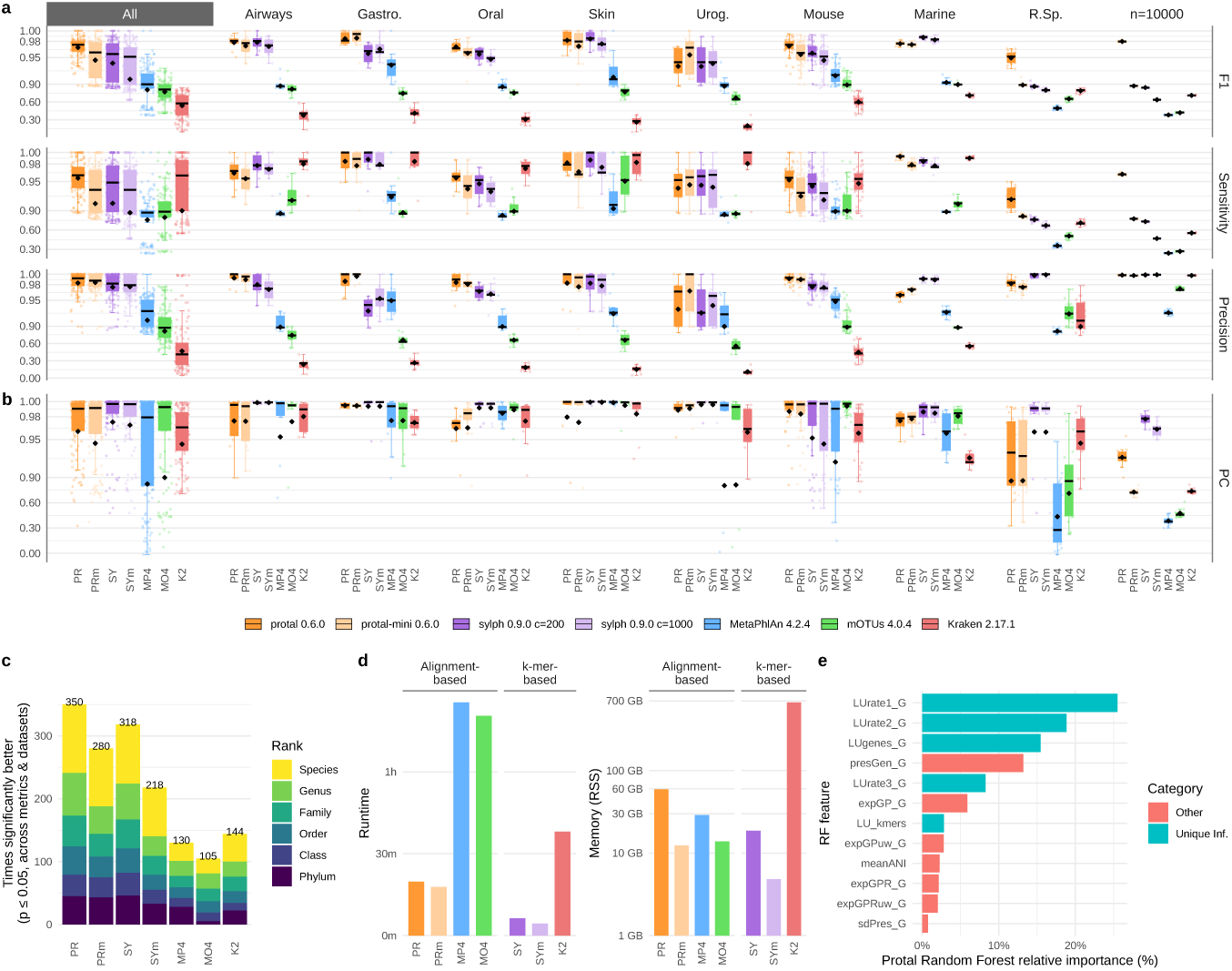
Comparative benchmarking of metagenomic taxonomic profilers. **a** Profiling accuracy across synthetic benchmark datasets, including CAMI environments (Airways, Gastrointestinal, Oral, Skin, Urogenital, Mouse and Marine), a rare-species benchmark (R.Sp.) and a high-complexity metagenome containing 10,000 species. F1-score, precision and sensitivity were calculated for each profiler; “All” summarises performance across all benchmark samples (see Suppl. Table S2. **b** Accuracy of species-abundance estimates, measured as the Pearson correlation between predicted and simulated species relative abundances. **c** Pairwise comparison of profiler performance at different taxonomic levels across the synthetic datasets shown in **a**. For each dataset and taxonomic level, paired t-tests were used to assess whether one profiler performed significantly better than another across samples (see Suppl. Table S3). **d** Runtime and peak memory use of each profiler, benchmarked on the same ten CAMI metagenomes, each containing 10 GB of read data, using 32 CPU cores. **e** Relative importance of internally computed per-taxon features used by the random forest classifier to predict taxon presence. Abbreviations: PR, protal; PRm, protal-mini; SY, sylph with c=200; SYm, sylph with c=1000; MP4, MetaPhlAn 4; MO4, mOTUs4; K2, Kraken2. Boxplots crossbars show medians, diamonds show means and whiskers show the Tukey range. Significances were determined via Wilcoxon rank-sum tests.

To resolve evolutionary relationships among strains detected in different metagenomes, protal reconstructs marker-gene-specific multiple sequence alignments (MSAs), for each species detected in at least three metagenomes (Fig. 1a). This step is implemented as a reference-guided MSA approach (RG-MSA), conceptually similar to ViralMSA [19], allowing strain-resolved metagenomic analyses to be performed with linear runtime and high computational efficiency.

### Reference databases

The species a reference-based profiler can detect, are limited by its underlying database. mOTUs4 [7] and MetaPhlAn 4 [6] build custom databases, using universal and species–specific marker genes, respectively. Instead, protal uses the standardized, externally curated GTDB taxonomy [5] (here, GTDB r226). GTDB was chosen as reference, that profiles stay compatible with a widely used genome taxonomy, and that future GTDB releases can be incorporated with minimal changes.

Built from GTDB r226, protal (and here Kraken2 and sylph) cover 143,614 species (29,405 genera and 5,932 families), while MetaPhlAn 4 and mOTUs4 represent half or fewer that numbers of species (Fig. 1b). This narrower coverage of the latter profilers partly reflects how species-specific marker databases are built: reliably identifying markers that are both unique to a species and conserved within it generally requires several genomes of that species, and MetaPhlAn 4, for example, defines a previously uncharacterized species only when at least five (metagenome assembled) genomes are available [6]. This is consequential because 65.5% of GTDB r226 species are represented by a single genome, which can be integrated in protals *universal* single-copy marker gene approach.

### Development and benchmarking of protal’s alignment algorithm

Protal uses a newly developed alignment algorithm based on exact k-mer matching with core-mers and flex-mers (Fig. 1c; Supplementary Fig. S5), which are compressed in memory using syncmers [15]. Candidate marker genes identified in this first step are then evaluated by a more precise, but computationally more expensive, wavefront alignment [16] (Methods; Supplementary Fig. S6).

To assess alignment performance in a metagenomic setting, we simulated reads from GTDB r214 genomes and aligned them against 120 species-representative marker genes using protal, Bowtie2 [23], BWA-MEM2 [24], minimap2 [25] and Strobealign [26]. Because the source genome of each simulated read was known, true-positive rates (TPR) and false-discovery rates (FDR) could be estimated across mapping-quality (MAPQ) thresholds.

Overall, most aligners showed similar sensitivity and precision, with Bowtie2 performing slightly worse (Fig. 1d). Protal achieved a TPR of 77.4% and an FDR of 3.0% at MAPQ 5, providing a balanced threshold for species-classification tasks; higher MAPQ thresholds reduced sensitivity with only limited gains in precision. Strobealign reached a higher TPR of 79.0% at MAPQ 60, but with a higher FDR of 3.7%. BWA-MEM2 showed slightly better accuracy than protal, with 78.0% TPR and 2.7% FDR at MAPQ 60, but required more memory and longer runtime. Bowtie2 had lower sensitivity, with 72.6% TPR at a comparable FDR of 3.2% using MAPQ 7.

Across five iterations, protal aligned 2 *×* 5 GB of input reads in 5 min 52 s, making it more than twofold faster than BWA-MEM2, more than threefold faster than Bowtie2 and more than fourfold faster than minimap2 (Fig. 1e). Strobealign was slightly faster than protal, completing the same task in 5 min 10 s, at slightly lower precision. The protal marker-gene index required 37 GB of memory, approximately twice that of Bowtie2 and minimap2, but less than BWA-MEM2 and slightly less than Strobealign (Fig. 1e).

Together, these results show that protal provides alignment performance comparable to established short-read aligners while producing additional taxonomically informative alignment features used for high-resolution metagenomic profiling.

### Protal detects species reliably and fast in metagenomes

We evaluated protal on nine synthetic metagenomic benchmark datasets designed to capture different profiling challenges. These comprised CAMI datasets [27, 28] from five human body sites (Airways, Gastrointestinal, Oral, Skin and Urogenital; *n* = 49 metagenomes in total), Mouse (*n* = 49) and Marine (*n* = 9) communities, together with a rare-species benchmark (R.Sp.; *n* = 20) and a high-complexity metagenomes containing approximately 10,000 species (*n* = 10, Fig. 2a). The R.Sp. dataset tested detection of species represented by only a single genome in GTDB r214 (reannotated to GTDB r226 taxonomy for benchmarking), whereas the high-complexity dataset tested robustness under extreme community complexity. Protal was compared with alignment-based profilers (MetaPhlAn 4 and mOTUs4) and k-mer-based profilers (sylph and Kraken2+Bracken) using a standardized GTDB-based taxonomy (Methods). GTDB delineates species by average nucleotide identity, which can resolve a traditionally recognised species into several genomically distinct ones — *Faecalibacterium prausnitzii*, for example, is represented by seven separate species clusters in our gold standard [5, 37]. Because the hardest calls to make are those between congeneric species, this finer delineation weighs mainly on precision, and species-level values are therefore not directly comparable with NCBI-taxonomy-based benchmarks.

Across all datasets, protal achieved the highest mean species-level F1-score (0.969*±* 0.021) and sensitivity (0.956 *±* 0.032), with a precision (0.983 *±* 0.026) matched only by protal-mini (0.984 *±* 0.021; Fig. 2a). Sylph was the strongest alternative overall (F1 = 0.939 *±* 0.050, all comparisons are available in Suppl. Table S2).

On standard CAMI human body-site and Mouse datasets, protal and sylph performed similarly: sylph achieved marginally higher sensitivity across body sites (0.969 versus 0.966), whereas protal retained higher precision (0.977 versus 0.958) and performed best on Mouse. The Marine dataset was the main exception, where sylph achieved higher F1-score and precision than protal (0.988 and 0.991 versus 0.976 and 0.959), while protal retained near-perfect sensitivity (0.993). Kraken2+Bracken had the lowest overall F1-score and precision, despite low-abundance filtering as recommended for Bracken [31].

Protal’s advantage was clearest in the more challenging simulations, indicating that protal retains species-level sensitivity and precision for both poorly represented and highly complex communities. In the rare-species benchmark, protal reached the highest F1-score (0.949 *±* 0.012) and sensitivity (0.919), outperforming sylph (F1 = 0.863 *±* 0.019; sensitivity 0.761), MetaPhlAn 4 (F1 = 0.495 *±* 0.025) and mOTUs4 (F1 = 0.653 *±* 0.028). In the high-complexity metagenomes, protal maintained high sensitivity (0.963*±*0.002) and precision (0.998*±*0.000), yielding an F1-score of 0.980*±* 0.001, compared with sylph (0.843), Kraken2+Bracken (0.711), MetaPhlAn 4 (0.382) and mOTUs4 (0.419).

This performance reflects two design choices: protal’s marker gene based profiling allowing to detect species represented by a single genome, and alignment-based lookups that are more sensitive than sketch matching. Analysis of the random-forest feature importances indicated that long unique k-mer gene rates (lu gene rate variants) were the strongest predictors of true taxon presence. Features more readily accessible via external aligners, such as gene-presence and alignment-quality (% identity, coverage variation), contributed considerably less to profiling performance (Fig. 2e), demonstrating the value for protal’s custom alignment algorithm approach.

We further evaluated reduced-memory configurations for protal and sylph. Protal-mini maintained near-identical precision to the full database (0.984) but reduced sensitivity, giving an overall F1-score of 0.945 *±* 0.041. Sylph with a sparser sketch (*c* = 1000; “sylph-mini”) showed lower overall performance (F1 = 0.909 *±* 0.098), particularly in the high-complexity metagenomes (F1-score 0.638).

Next, we tested how often one profiler’s F1-score, sensitivity or precision was significantly better than another’s, across the nine datasets and all taxonomic levels from phylum to species (paired t-test, Fig. 2c). Here, protal excelled, being in 350 comparisons better than other profilers, followed by sylph, protal-mini and sylph-mini (318, 280, 218 times better in comparisons).

### Analysis of taxonomic misassignment

For every profiler at least 85% of false-positive calls — 89% for protal, essentially all for sylph and MetaPhlAn 4 — were congeneric with a species in the gold standard of the same sample, and most of these species were below 0.1% relative abundance. Profilers differ in how many misassignments they make: protal and protal-mini produced fewer than 500 each across all 137 benchmarking samples, against more than 22,000 for Kraken2+Bracken.

That difference was large, and it was clearest in species-poor communities, where a given number of false calls represents a larger fraction of the community. Ranking the profilers by false positives per 100 true species (across the five human body-site datasets, averaging 31–77 species per sample) gives protal-mini (1.5), protal (1.7), sylph (3.3 at both sketch densities), MetaPhlAn 4 (8.6), mOTUs4 (42) and Kraken2+Bracken (442). The same broad ordering held across all nine datasets (medians 1.8, 1.4, 1.6–2.4, 6.9, 28.6 and 279 false positives per 100 true species, respectively).

For the two lowest-ranked tools the number of false calls per sample stayed high even in the smallest communities, so their precision was governed mainly by community size: mOTUs4 ranged from 0.56 in the species-poor Urogenital samples (31 species) to 0.97 in the high-complexity metagenomes (10,000 species), and Kraken2+Bracken from 0.11 to 0.998 over the same range. Precision measured on species-rich communities alone therefore substantially overstates how these tools behave on the small communities typical of many host-associated body sites.

### Accurate compositional and diversity reconstructions across community types

We next evaluated whether profilers accurately reconstructed species abundances and community richness. Protal showed high agreement between predicted and expected species abundances, outperforming Kraken2+Bracken, MetaPhlAn 4 and mOTUs4, with a mean Pearson correlation of 0.961 *±* 0.083 at species level (Fig. 2b); similar trends were observed at genus level (Suppl. Fig. S1). Sylph achieved marginally higher abundance correlation (0.973 *±* 0.077).

We assessed richness-dependent performance using the 49 synthetic CAMI Mouse metagenomes, which span 47–223 species in the gold standard (Suppl. Fig. S2). Predicted richness closely matched the expected richness for protal, sylph and MetaPhlAn 4, with average deviations of approximately 4–7%. In contrast, Kraken2+Bracken overestimated richness by roughly twofold, predicting on average 295 species compared with 143 in the gold standard. Among correctly identified species, protal recovered true positives at relative abundances as low as 0.0004% in the CAMI Mouse data, comparable to mOTUs4 (0.0005%) and well below the detection floor of MetaPhlAn 4 (0.0012%) and sylph (0.0023%). Kraken2+Bracken reached a nominally lower value (0.0001%), but only as a by-product of its indiscriminate over-prediction, consistent with its roughly twofold richness inflation and elevated false-positive rate.

Profiling performance remained robust across this richness gradient. Whereas the comparison above contrasted whole datasets spanning 31 to 10,000 species, this analysis follows the same profilers across the narrower richness gradient within a single dataset. Although false negatives increased with richness for all profilers (*p <* 0.001), sensitivity declined only weakly and non-significantly. The main difference between profilers was the behaviour of false-positive calls: false positives increased significantly with richness for sylph, MetaPhlAn 4 and protal-mini (*R* = 0.54–0.70, all *p <* 0.001), and more weakly for mOTUs4 (*R* = 0.40, *p <* 0.01) and Kraken2+Bracken (*R* = 0.30, *p <* 0.05), whereas protal was the only profiler for which false-positive counts were independent of richness (*R* = 0.16, ns). Thus, protal preserved precision as community complexity increased, also reflected in its profiling performance of high complexity community (Fig. 2b)

### Computational performance

Profiler runtime was compared using ten CAMI Airways metagenomes processed under identical settings (Fig. 2d). As expected, the k-mer-based profilers were fastest, with sylph completing the benchmark in 6 min 18 s (*c* = 200; 4 min 25 s at *c* = 1000) and Kraken2+Bracken in 38 min. Among alignment-based profilers, protal was fastest, completing the benchmark in 19 min 45 s, compared with 1 h 25 min for MetaPhlAn 4 and 1 h 21 min for mOTUs4. Thus, protal reduced alignment-based profiling time by 4.3-and 4.1-fold relative to MetaPhlAn 4 and mOTUs4, respectively, while protal-mini was faster still (17 min 52 s) with a five-fold memory reduction (12 GB versus 59 GB).

Although metagenomic sequence alignment increases computational cost compared with k-mer-based profiling, it provides read-level information that enables downstream *de novo* reconstruction of strain-resolved phylogenies, as evaluated in the next section.

### Protal reliably resolves strains at species-profiling runtime

Protal resolves intraspecific phylogenies by constructing one reference-guided marker-gene MSA per detected species, followed by a phylogenetic reconstruction with IQ-TREE. We benchmarked this functionality using STRAIN46, a synthetic dataset of 200 metagenomes containing 46 bacterial species, excluding conspecific strains within each metagenome (Methods). We compared protal with MetaPhlAn 4+StrainPhlAn 4 [6]; sylph and Kraken2 were excluded because they do not reconstruct intraspecific phylogenies *de novo*, and mOTUs4/metaSNV was excluded because the workflow could not be executed in our benchmark environment (Methods). Alongside protal, we also benchmarked protal-mini, to test the performance of a compact index.

Overall, runtime was lower for protal: its reference-guided MSAs added minor computational cost beyond species profiling. Across STRAIN46 subsets of 4–128 samples, protal was 6–40-fold faster end-to-end than MetaPhlAn 4+StrainPhlAn 4 (Fig. 3a). For example, at 64 samples protal completed profiling and strain resolution in 22 min, compared with 7.9 h for MetaPhlAn 4+StrainPhlAn 4. Protal required the same peak memory as for species level profiling (59 GB versus 29 GB MetaPhlAn 4+Strain-PhlAn 4), The strain-resolution steps themselves accounted for only 24–39% of protal’s total runtime (9.8 min of 41.1 min at 128 samples), whereas the strain-specific steps of the MetaPhlAn 4+StrainPhlAn 4 workflow dominated its runtime (43–82%).

**Fig. 3.**
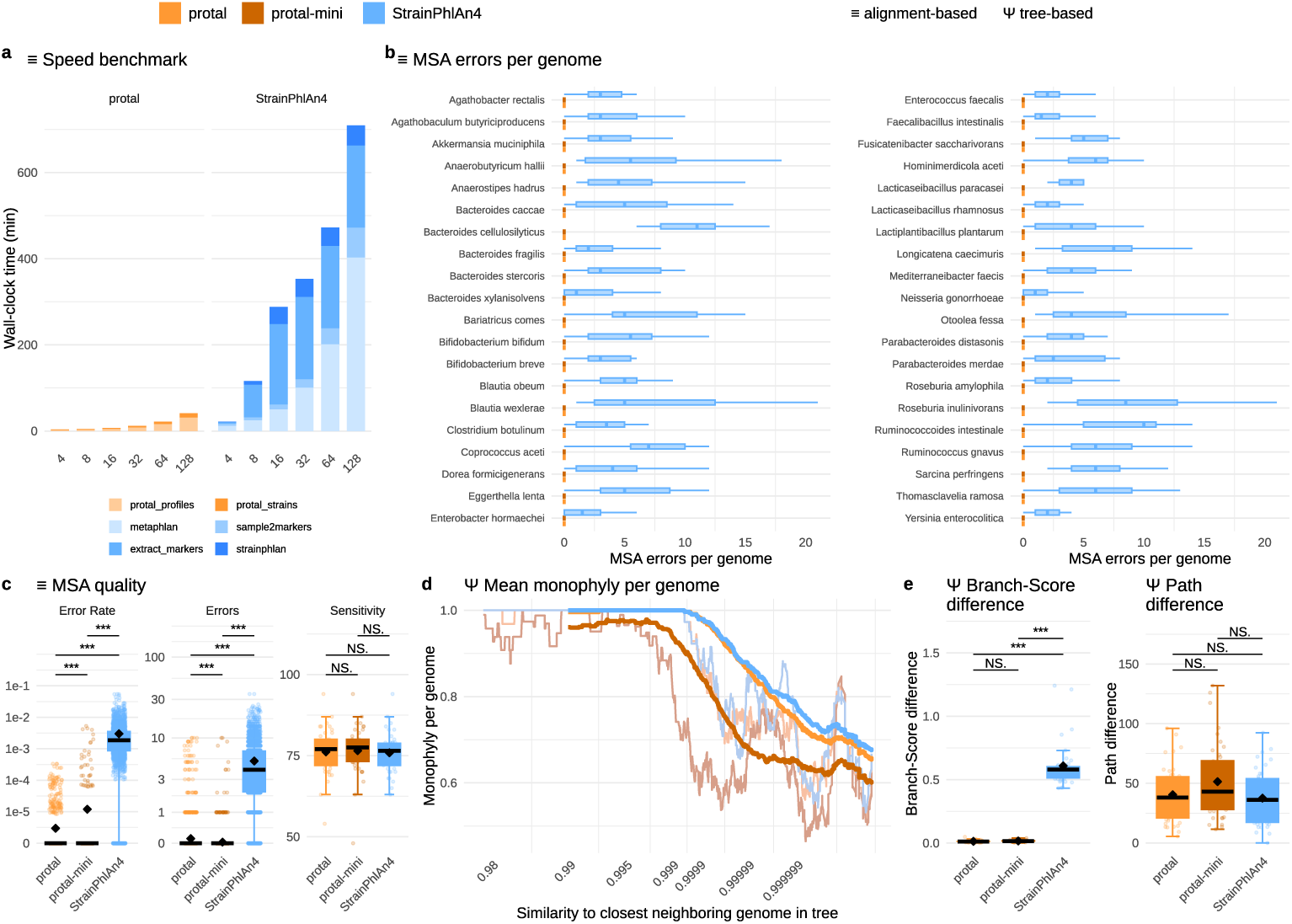
Protal enables strain-resolved profiling with reduced runtime and phylogenetic accuracy comparable to StrainPhlAn 4. **a** Runtime of protal and MetaPhlAn 4+StrainPhlAn 4 for increasing numbers of samples (*N* = 4–128) from the synthetic STRAIN46 dataset, stacked by pipeline stage. No species-or strain-level filtering was applied, phylogeny reconstruction runtime is not included. **b** Multiple-sequence-alignment (MSA) errors, measured as the number of miscalled sites relative to the known genome sequence, separated by species and tool. **c** MSA errors summarized as (left to right) fractional error rates, total error count (points represent species), and resolved species (points represent metagenomes). **d** Fraction of strains from the same simulated genome correctly resolved into monophyletic clades, compared to each strain’s similarity to its closest neighbouring genome (average nucleotide identity). Averages are visualized with a right-aligned sliding window over 250-genomes (coloured lines) and 30-genomes (grey). **e** Agreement between inferred *de novo* strain phylogenies and the gold-standard reference phylogeny. Inferred and reference phylogenies were restricted to shared tips (*≥* 4 shared tips), and similarity was quantified using branch-score distance [35] and path difference [36]. Boxplot marks are as in Fig. 2.

### Comparison of alignments and phylogenetic resolution

We first compared the marker-gene MSAs produced by the workflows (Fig. 3c). Only species not filtered by StrainPhlAn 4 or protal for technical reasons are shown (40 of 46 total, one and 5 filtered, respectively, for technical reasons).

Protal recovered the large majority of genomes without a single miscalled site: 95.3% of per-genome alignments were error-free (mean 0.13, median 0 errors per genome), compared with 7.8% error-free for StrainPhlAn 4 (mean 5.1, median 4; Wilcoxon rank-sum test, *p <* 0.001). Protal retains constant alignment positions and produces accordingly an approximately 30-fold longer alignment (median 121 kb versus 3.9 kb per species, respectively), which results in an error rate of 2.7*×*10*^−^*^6^ *±*1.9*×*10*^−^*^5^ protal versus 3.0*×*10*^−^*^3^ *±*4.0*×*10*^−^*^3^ StrainPhlAn 4. Protal-mini, built on a much smaller marker set (median MSA length 16 kb), had 98.7% error-free genomes and on average 0.03 errors per genome. Species retention was comparable among tools (median 76 *−* 77, Fig. 3c).

This advantage was consistent across the species panel rather than driven by a few outliers (Fig. 3b). Stratifying per-genome MSA errors by species, protal and protal mini had a median of zero errors per genome, whereas StrainPhlAn 4 had a non-zero median in all reconstructed species (1–11 errors per genome; median= 4). Strain-PhlAn 4’s noisiest reconstructions were those for *Bacteroides cellulosilyticus* (median 11 errors per genome, up to 24) and *Ruminococcoides intestinale* (median 10, up to 22), for both of which protal produced error-free alignments for every genome.

We next assessed strain resolution by testing whether genomes seeded from the same strain were recovered as monophyletic clades[38], on the same 40 species as above. As expected, resolution decreased with more similar genomes: both tools resolved strains reliably *<* 99.9% nearest-neighbour similarity, but correct resolved monophyly declined to *∼* 65% at *>* 99.9999% (Fig. 3d). Overall monophyly was comparable between protal and StrainPhlAn 4 (0.867 *±* 0.245 versus 0.877 *±* 0.236; *p* = 0.38), whereas protal-mini, with its smaller marker set, resolved strains less reliably (0.793 *±* 0.307; *p ≪* 0.001).

### Resolving phylogenetic relatedness

For the same 40 species, we compared inferred intraspecific phylogenies with gold-standard whole-genome phylogenies built with Roary [32]. Because protal retains constant (invariant) alignment positions by default, its branch lengths are expressed on a per-site scale directly comparable to the whole-genome reference.

Summed over shared tips, protal’s phylogenies were 0.65-fold the length of the gold-standard phylogeny (median; IQR 0.52–0.79), consistent with marker genes being modestly more conserved than the genome average. In contrast, StrainPhlAn 4’s alignments of only variable-site inflated phylogenies’ length roughly 20-fold (median ratio 21.6; IQR 16.6–34.2). Consequently, protal’s phylogenies were about 50-fold closer to the gold standard in branch-score distance [35] than StrainPhlAn 4’s (median 0.012 versus 0.580; lower in 40 of 40 shared species; *p <* 0.001, Wilcoxon rank-sum test), and protal’s phylogeny lengths’ tracked the reference more closely across species (Pearson *R* = 0.78 versus 0.64). However, phylogeny topology was recovered equally well by both workflows, with no significant difference in path difference [36] (median 38.1 versus 36.1; *p* = 0.59; Fig. 3e).

## Discussion

Protal demonstrates that alignment-based metagenomic profiling can be made sufficiently fast for large-scale studies while retaining the nucleotide information required for strain-resolved analysis. Across synthetic benchmarks, protal combined complete GTDB species coverage, high species-level precision and sensitivity, accurate abundance reconstruction, and substantially faster runtime than existing alignment-based profilers. Unlike purely k-mer-based methods, protal also reconstructs marker-gene alignments for detected species, enabling *de novo* inference of intraspecific phylogenies within the same workflow.

### Algorithmic design and reference strategy

Protal’s performance is driven by a purpose-built alignment strategy based on core-mers and flex-mers stored in a custom hash-based data structure. This design is tailored to large collections of related marker-gene sequences, where many regions are conserved but only a subset of positions is taxonomically informative. By storing additional information, including k-mer uniqueness and taxonomic informativeness, protal can use alignment-derived features for downstream species prediction rather than relying only on best-hit alignments or exact k-mer matches. The use of unique k-mers was inspired by KrakenUniq [39], which implements a counting of unique k-mers to improve precision, and GT-Pro [40], which pre-selects informative k-mers, placing protal conceptually between conventional short-read aligners (usually FM-index-based) and sketch-based taxonomic profilers. The former while generalizable and memory efficient, are not optimized for profiling against highly-redundant marker-gene databases. Sketch-based profilers (e.g. Kraken2[8] and sylph[9] benchmarked here), achieve high speed by reducing the search space, but generally do not retain the alignments needed to reconstruct strain-level sequence variation [8, 9]. Protal instead combines fast hash-based lookup and alignment validation to support taxon validation, abundance estimation and strain-resolved phylogenetics.

A second key design choice is direct integration with GTDB [5]. This provides compatibility with a standardized, externally curated taxonomy and allows protal databases to be rebuilt as GTDB expands [37], such as with the recent long-read based environmental MAGs [41]. Because protal uses universal marker genes from GTDB representative genomes, it can represent bacterial species even when only a single genome is available. This substantially increases taxonomic coverage compared with profilers that rely on custom marker databases requiring broader within-species genome representation, such as MetaPhlAn 4 [6].

### Marker-gene choice creates a trade-off between breadth and strain resolution

Protal uses 120 universal bacterial and 53 archaeal single-copy genes, providing broad species coverage and a standardized basis for both taxonomic profiling and strain-resolved analysis. Different subsets of universally conserved genes have been used for metagenomic species profiling and genome-based taxonomic frameworks [42, 43, 5], due to being taxonomically stable within the phylogenetic history of a species and often exempt from horizontal gene transfers [14, 13].

However, the species-specific markers individually selected for each species and used by StrainPhlAn 4 contain more intraspecific variation due to being faster-evolving [44]. This may a) provide a higher fidelity to identify a species correctly (e.g. MetaPhlAn 4 versus mOTUs4 across all datasets, mean species-level F1 0.80 versus 0.77 and precision 0.91 versus 0.81; Fig. 2a; this ordering reverses on the rare-species and high-complexity simulations, where reference breadth rather than marker resolution limits detection) and b) provide a higher intraspecific strain-resolution. Consistent with a marker-resolution limit, species-level errors in every profiler were dominated by misassignments among close relatives rather than detection of absent lineages; what differed was their frequency, with mOTUs4 making about five times more such calls than MetaPhlAn 4 in the species-poor body-site communities (42 versus 8.6 false positives per 100 true species).

Conversely, protal’s conserved marker genes may provide a more stable signal across broader evolutionary distances, at the expected cost of resolving near-identical strains. In our benchmark this trade-off was smaller than expected: overall strain resolution did not differ significantly between the two tools (*p* = 0.38), and only above 99.99% nearest-neighbour similarity did StrainPhlAn 4’s faster-evolving markers hold a slight edge (mean monophyly 0.726 versus 0.702), while between 99.9 and 99.99% protal was ahead (0.936 versus 0.927). Further, protal can use information collected during alignment to improve species-detection confidence. Thus, the two tools reach equivalent strain-tree topologies through different marker-gene strategies, while protal additionally recovers cleaner alignments and branch lengths on the correct absolute scale.

### Relationship to existing profiling and strain-resolved approaches

K-mer-based profilers are extremely fast and can identify taxa or strains represented in their databases, but they are less suited to reconstructing genetic distances among previously unobserved strains. Kraken2 can classify reads to known strain labels when such references are present [8], and GT-Pro efficiently detects predefined SNPs [40], but neither reconstructs *de novo* strain phylogenies from metagenomes. Sylph showed excellent species-profiling performance in our benchmarks, but its sketch-based design does not retain nucleotide alignments for strain-sequence reconstruction [9].

Alignment-based approaches are therefore better suited for discovering and comparing unknown strain variation, but have historically been computationally expensive. StrainPhlAn provides strain-level population structure from metagenomes, but requires a dedicated downstream workflow after species profiling [11, 6]. Protal addresses this limitation by making alignment-based profiling substantially faster while retaining sufficient sequence information for reference-guided MSA construction and phylogenetic inference. The two approaches also differ in what their trees can be used for: because protal retains constant alignment positions, its trees carry branch lengths on the same per-site scale as a whole-genome phylogeny, whereas variablesite-only alignments inflate branch lengths by roughly an order of magnitude. This matters wherever absolute divergence is interpreted — dating transmission, comparing evolutionary rates across species, or pooling trees across studies — rather than only tree topology. For samples dominated by taxa that are poorly represented in reference databases, assembly-or MAG-based approaches remain preferable because they can recover larger genome fractions and functional content [38, 45, 46].

### Limitations and outlook

Protal has several current limitations, most of which reflect deliberate design choices rather than fundamental constraints. First, protal does not currently deconvolute multiple conspecific strains within a single metagenome. Instead, strain-resolved analyses are restricted to cases with a dominant strain signal, an approach commonly used in strain-resolved metagenomics [11, 38]. Fully *de novo* resolution of co-occurring conspecific strains remains computationally demanding [47], whereas faster alternatives typically depend on strain-level references or predefined variant panels [48, 49, 50]. Such reference dependence would conflict with protal’s aim of reconstructing strain relationships without requiring prior knowledge of the strains present.

Second, species detection depends on the quality and structure of the reference taxonomy. Chimeric reference genomes, imprecise species boundaries, horizontal gene transfer and homologous recombination can all reduce specificity by blurring the genetic distinction between closely related taxa.

Third, because protal relies on universal marker genes, strain comparisons are based on approximately 5% of the genome. This supports broad taxonomic coverage and computational efficiency, but limits resolution for strains separated by very few substitutions, and means that detection ultimately depends on coverage of a small genomic fraction; protal nonetheless recovered true positives down to 0.0004% relative abundance, below the detection floor of the other marker-gene-based profilers tested. Protal is also currently restricted to prokaryotic species and short-read metagenomic data. Extension to eukaryotes will require separate marker-gene selection, database construction and validation, as eukaryotic genome representation, gene structure and marker-gene behaviour differ substantially from those of bacteria and archaea. Future versions will also support long-read metagenomic data, where longer alignments may improve both species assignment and strain reconstruction.

Finally, reduced-reference configurations provide a practical route to lower memory requirements. A protal-mini database using 20 marker genes reduces the memory footprint roughly fivefold (12 GB versus 59 GB peak resident memory) with only a modest reduction in profiling accuracy (F1 0.945 versus 0.969), bringing species profiling within reach of laptop-class hardware. Strain resolution is more affected: protal-mini recovered fewer strains as monophyletic (0.793 *±* 0.307 versus 0.867 *±* 0.245), with the loss concentrated among near-identical genomes (mean monophyly 0.671 versus 0.936 between 99.9 and 99.99% nearest-neighbour similarity). The reduced database is therefore best suited to species profiling and to strain analyses of well-separated lineages, while the full database remains preferable for fine-grained strain resolution.

The advances integrated in protal will enable laboratories with different levels of computational infrastructure and expertise access to fast and reliable, alignment-based and strain-resolved metagenomic profiling, enabling a new class of studies.

## Availability

protal is available as open source software under a MIT license from https://github.com/4less/protal/ and can be installed from https://anaconda.org/bioconda/protal. The reference database build on GTDBr214 is available at https://protal.quadram.ac.uk.

## Funding

JF was supported by the UKRI Biotechnology and Biological Sciences Research Council Norwich Research Park Biosciences Doctoral Training Partnership, BB/T008717/1. FH was supported by European Research Council H2020 StG (erc-stg-948 219, EPYC). FH and AD gratefully acknowledge the support of the Biotechnology and Biological Sciences Research Council (BBSRC); this research was funded by the BBSRC Institute Strategic Programme Food Microbiome and Health BB/X011054/1 and its constituent project(s) BBS/E/QU/230001B (Theme 2, The impact of plant-based foods on the GI microbiome, keystone species and food-borne pathogens) and Biotechnology and Biological Sciences Research Council (BBSRC) Institute Strategic Programme (ISP) BB/X011054/1 and its constituent project BBS/E/F/000PR13631; Earlham ISP BBX011089/1 and its constituent work package BBS/E/ER/230002A.

## Acknowledgments

We would like to acknowledge the input of the entire Hildebrand group at the Quadram Institute. Furthermore, we would like to thank the Earlham Institute HPC and Research Computing teams for their continued support. ChatGPT 5.5 was used to revise text clarity.

## Author contributions

Conceptualization - JF, FH; Data curation - JF; Formal analysis - JF; Funding acquisition - FH; Investigation - JF, FH; Methodology - JF, FH; Project administration - FH; Resources - JF, FH; Software - JF; Supervision - FH; Validation - JF, AD; Visualization - JF, FH; Writing – original draft - JF, FH; Writing – review & editing - FH, JF, AD

## Supplementary Figures

**Fig. S1.**
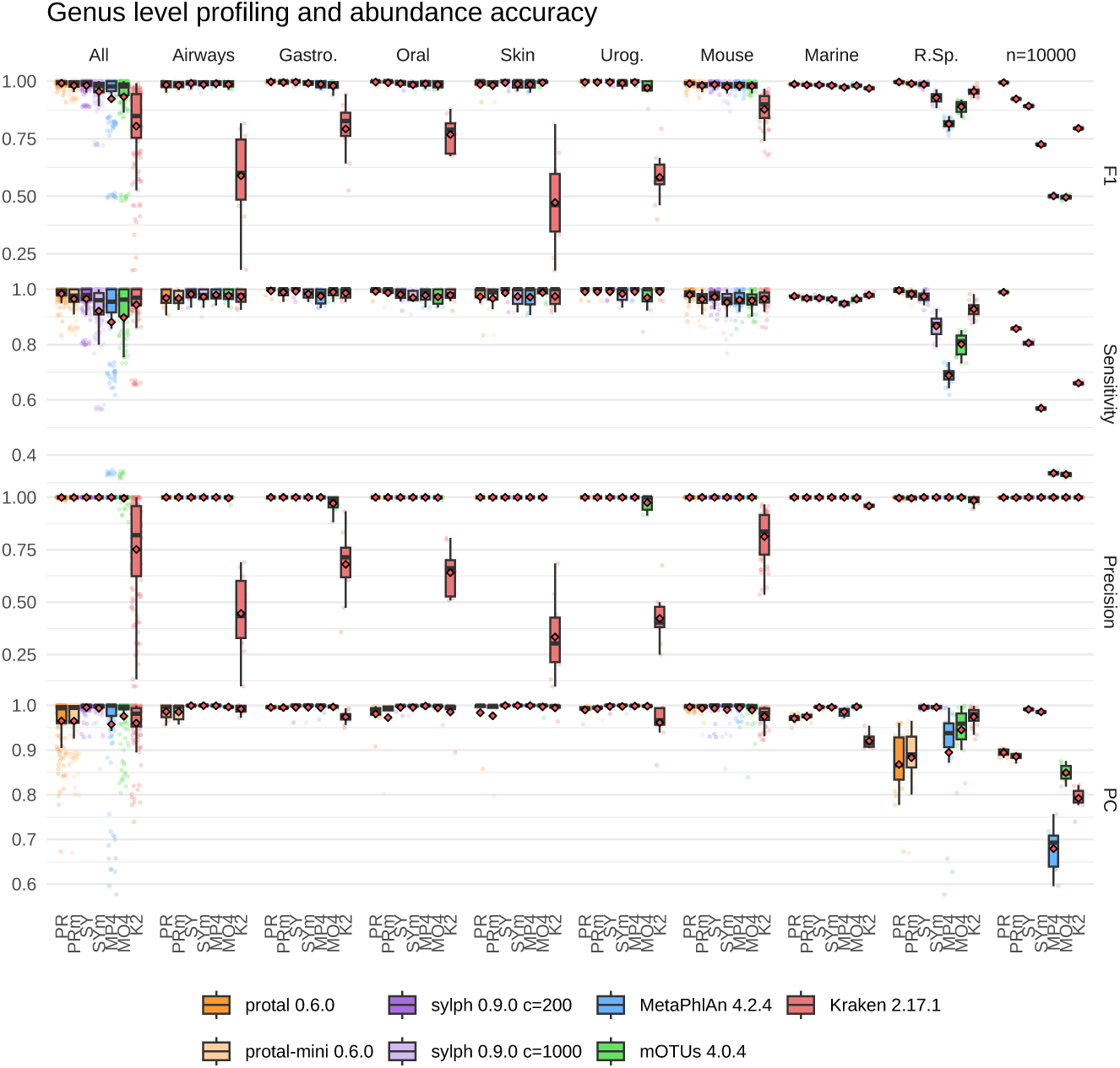
Profiling and abundance benchmark for taxonomic rank Genus, across the same benchmark datasets as Fig. 2. Rows show F1-score, sensitivity and precision, and abundance accuracy measured as the Pearson correlation (PC) between predicted and simulated genus relative abundances. Box-plots show the distribution across samples, with red diamonds marking the mean. “All” summarises performance across all benchmark samples.

**Fig. S2.**
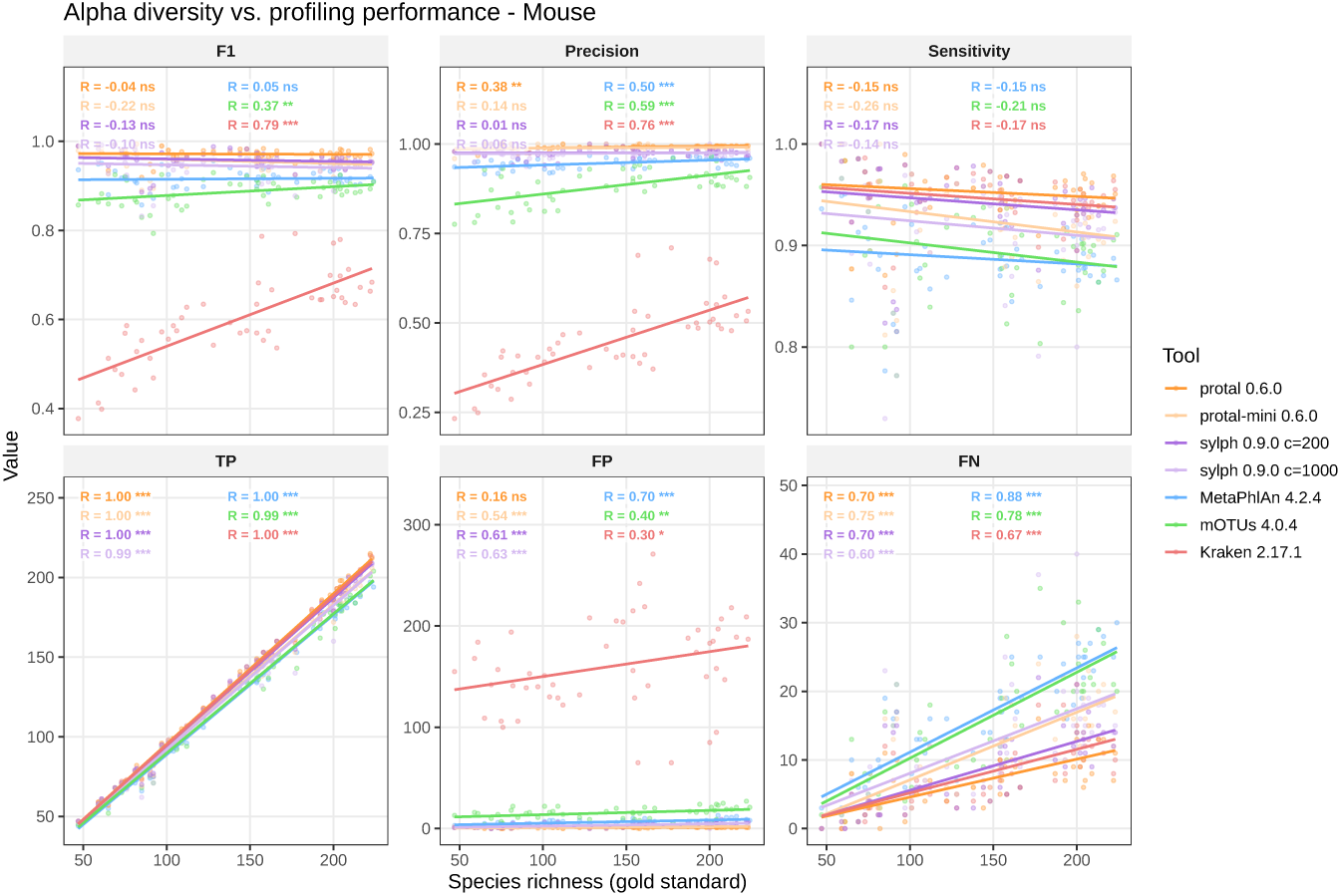
Profiling performance with respect to sample alpha diversity (species richness) in the CAMI Mouse dataset. Each dot represents one sample and all statistics are after adjustment on species level. Species richness on the x-axis is the gold-standard number of species (TP+FN) and each panel shows the respective metric on the y-axis: F1-score, precision, sensitivity, true positives (TP), false positives (FP), and false negatives (FN). Lines are fitted per tool and annotated with the Pearson correlation coefficient (R) and significance (ns: not significant, *: p¡0.05, **: p¡0.01, ***: p¡0.001).

**Fig. S3.**
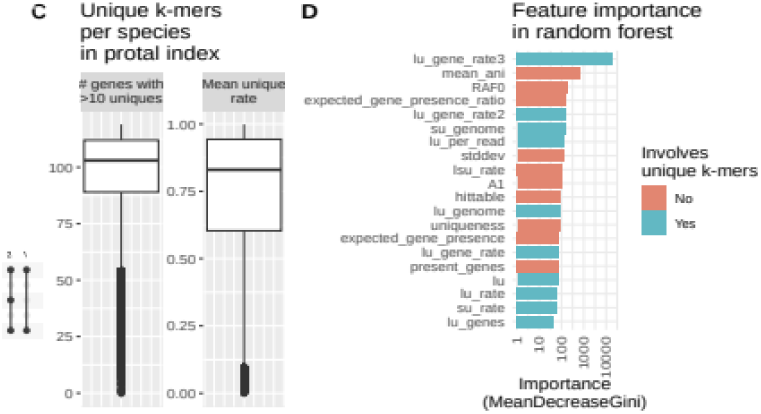
Key features to increase species detection. a The number of genes containing at least 10 unique k-mers (short uniques and long uniques) across all species, being on median 103 genes. However, some species contain significantly less marker genes, thus being harder to uniquely identify. b Variable importance reported from constructing the random forest. MeanDecreaseGini quantifies the importance of a variable within the random forest.

**Fig. S4.**
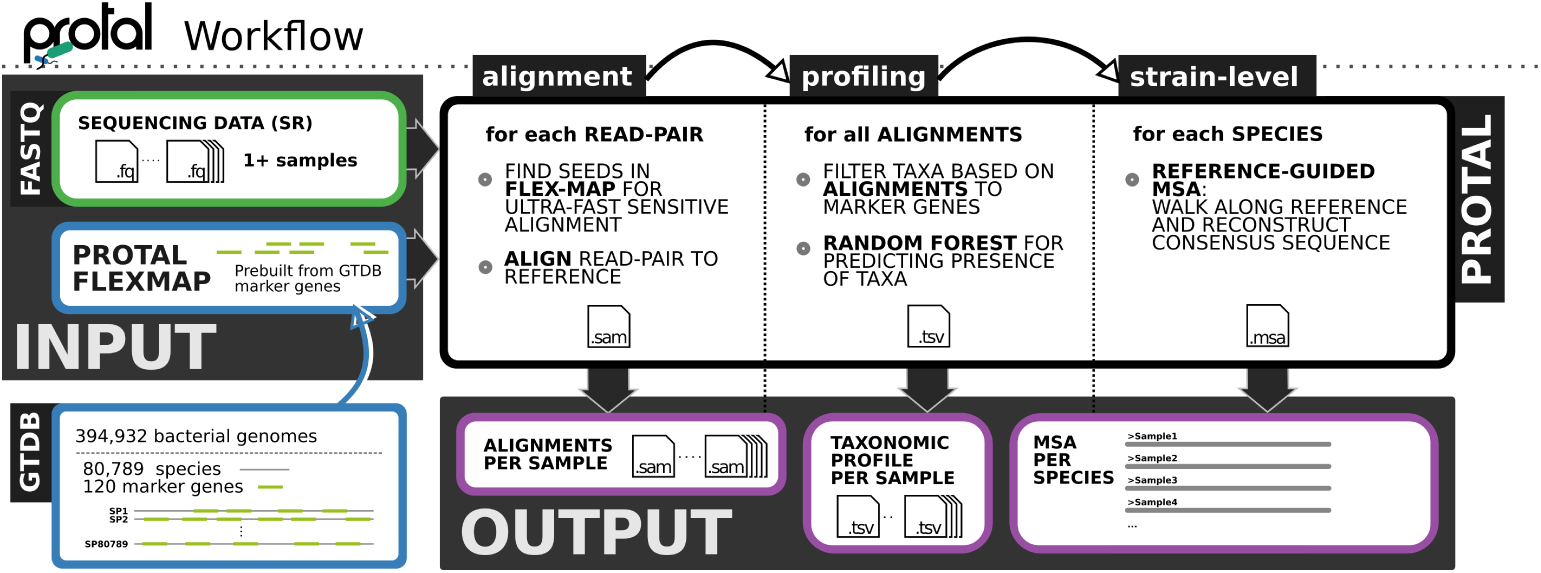
Protal takes a set of paired-end short reads from shotgun metagenomic sequencing and outputs per sample taxonomic profiles and strain-resolved MSAs per species present in multiple samples. Internally, protal has three distinct steps - alignment, profiling, and strain-level. In the first step, all reads are aligned against all species-representative marker genomes within GTDB. During profiling, these alignments are processed and counted for each marker gene of each species. A random forest evaluates evidence for each species to predict presence or absence. To achieve strain-level resolution, alignments from the same species across multiple samples are used to yield a reference-guided alignment.

**Fig. S5.**
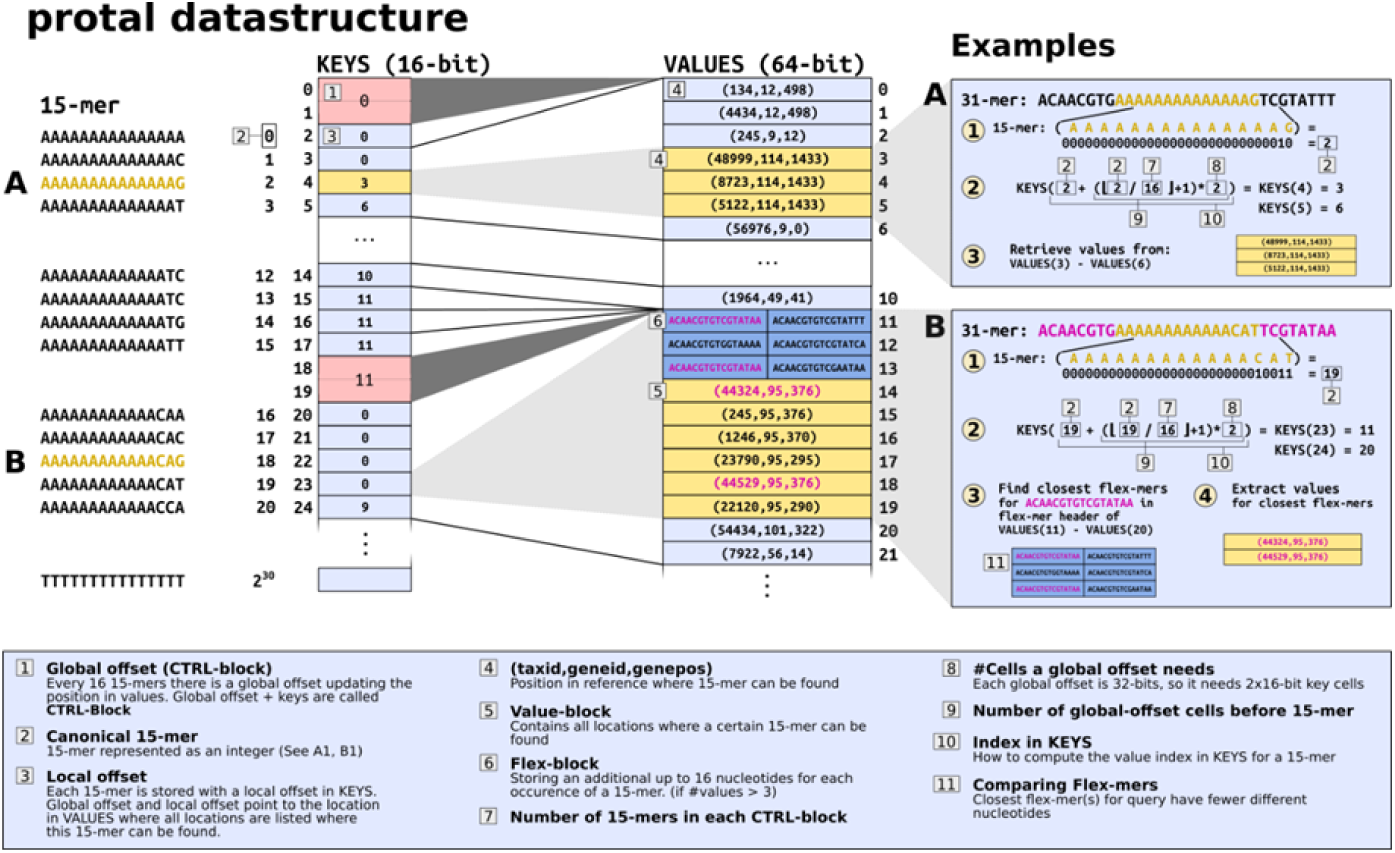
The data-structure that holds the index build from a set of reference sequences with known taxonomic and genic origin. The data-structure maps core-mers (KEYS) with all their occurrences (VALUES) in the provided reference sequences. To perform a lookup, the integer (canonical) representation of a k-mer is used to compute the index in the KEYS table. The index is the absolute integer value of the k-mer plus the number of preceding cells that belong to CTRL Blocks. The closest preceding CTRL block value plus the offset number at the KEYS cell gives the index for the VALUES table that contains the corresponding 3-tuples pointing to locations in the reference sequences as shown in a2 and b2. If a core-mer has more than 3 occurrences, flex-mers are used to find the closest matching reference. Example a and b show how querying without, and with flex-mers works, respectively.

**Fig. S6.**
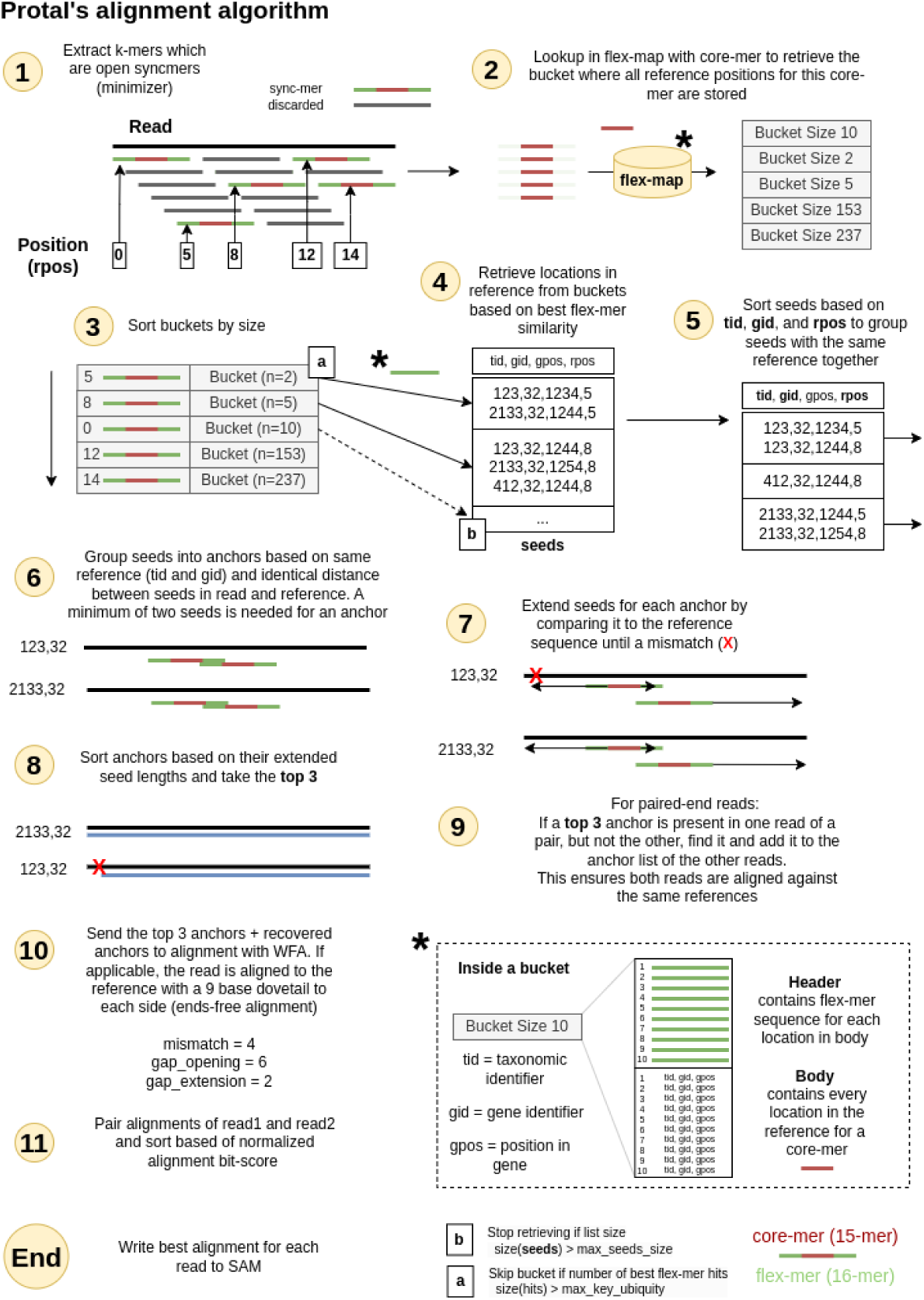
Detailed description of protal’s alignment workflow. See Section 3 for more details. Information about unique k-mers is not mentioned here, as they do not influence the alignment process.

## Supplementary Table Legends

**Suppl. Table S1** Alignment-derived per-species metrics used by the random-forest classifier to predict species presence in a metagenomic sample. The Short label column gives the abbreviations used for the features shown in the random-forest variable-importance panel (Fig. 2e); a trailing G marks metrics that are computed per gene and then summarized across genes.

**Suppl. Table S2** F1, precision and sensitivity of protal, sylph, MetaPhlAn 4, mOTUs4 and Kraken2, as shown in Fig. 2.

**Suppl. Table S3** Pairwise comparisons of profiler performance at different taxonomic levels

## Methods

### Reference database construction

Protal builds reference databases from GTDB species-representative genomes. Unless stated otherwise, analyses in this study used GTDB r226 [5] and the GTDB marker-gene set, comprising up to 120 bacterial and 53 archaeal universal single-copy marker genes per species. Marker genes were identified using the TIGRFAM [14] and PFAM [13] protein families used by GTDB. The default database used for species profiling contained 143,614 GTDB r226 species.

For reduced-memory analyses, we additionally generated a protal-mini database containing a reduced set of 20 marker genes.

### Specialized k-mer structure and index generation for fast and sensitive sequence lookup

#### Core-mers, flex-mers and the flex-map

Aligning short reads against large marker-gene databases requires an index that balances query speed, sensitivity and memory use. Protal uses a seeding strategy based on two complementary k-mer representations: core-mers and flex-mers. A core-mer is the central 15-mer of a 31-mer and is used for exact lookup. A flex-mer is the 16-nt sequence formed by the two 8-nt flanks surrounding the core-mer and is used to distinguish candidate hits with similar or identical core-mers (Fig. 1c; Supplementary Fig. S5). During lookup, protal first matches the 15-mer core exactly and then ranks candidate reference locations according to the similarity of their flanking flex-mers. The 15-mer core length was chosen because short exact seeds of similar size provide a useful sensitivity–specificity trade-off in established aligners such as minimap2 [25].

The protal index, termed the flex-map, stores locations of marker-gene k-mers in two arrays: a fixed-size 16-bit address array, KEYS, and a variable-size 64-bit value array, VALUES. The 30-bit two-bit encoding of a 15-mer core-mer, with A, C, G and T encoded as 00, 01, 10 and 11, respectively, is used as the direct address into KEYS. For example, the core-mer AAAAAAAAAAAAAGT maps to the bit string 000000000000000000000000001011, corresponding to index 11.

Each 64-bit cell in VALUES stores a hit tuple (tid, gid, gpos, unique), specifying the taxon identifier, marker-gene identifier, position within the marker gene and a 2-bit uniqueness flag. A value-block is the contiguous set of VALUES entries associated with the same core-mer. Core-mers absent from the reference have no corresponding value-block. If a core-mer occurs more than once in the reference, protal additionally stores or evaluates the associated 16-nt flex-mer to reduce the number of candidate reference locations returned for subsequent alignment.

To reduce the memory footprint of the address array, KEYS uses 16-bit local offsets into VALUES combined with periodic 32-bit control blocks. Entries are grouped in blocks of eight. Each block contains eight 16-bit local offsets and one 32-bit global offset, stored as two additional 16-bit cells. The local offsets in the block are interpreted relative to the block-specific global offset. This yields a constant KEYS size of 2^30^ *·* 2, bytes + (2^30^*/*8) *·* 4, bytes = 2.5, GB.

A naive implementation using one 64-bit offset per possible 15-mer would require 2^30^ *·* 8bytes = 8GB. The size of VALUES depends on the number of indexed marker-gene locations.

#### Unique k-mers

In addition to accelerating alignment, the flex-map stores information about the taxonomic uniqueness of k-mers, as unique k-mers provide evidence for reliable species detection. Protal annotates three classes of unique k-mers across the GTDB reference collection (Supplementary Fig. S3a). Short-unique k-mers are 15-mer core-mers found only within a single species. Long-unique k-mers are full 31-mers, comprising the core-mer and both flanking regions, with no occurrence outside their respective species. Long-super-unique (LSU) k-mers are long-unique k-mers whose closest non-conspecific match has Hamming distance of at least two; each LSU k-mer is therefore also a long-unique k-mer.

Uniqueness information is inferred during database construction by comparing stored k-mers against all GTDB r214 reference genomes. For the r226 database, this comprised 715,230 bacterial and 17,245 archaeal GTDB genomes [37]. Protal stores uniqueness information in the 2-bit uniqueness field of each VALUES entry. This information does not alter the alignment procedure itself, but is reported with the best alignment and used as input to downstream random-forest-based taxon detection.

#### Seeding, anchor finding and alignment

Protal aligns read pairs in three stages: seeding against the flex-map, grouping compatible seeds into anchors and extending the top-ranked anchors into full gapped alignments.

A **hit** is a reference location stored in VALUES as (tid, gid, gpos, unique). A **value-block** is the set of hits sharing the same core-mer. A **seed** is a query–reference match represented as (tid, gid, gpos, rpos, unique), where rpos is the position of the core-mer in the read.

For each read, protal extracts 31-mers that satisfy an open-syncmer criterion (Fig. 1c). Syncmers provide a more even subsampling of sequence space than minimizers [15]. The core-mer of each selected 31-mer is used as a key into the flex-map, and the corresponding value-block is retrieved from VALUES. Value-blocks are sorted by size in ascending order, so that more informative, less repetitive seeds are evaluated first. Starting with the smallest value-blocks, protal retrieves seeds and ranks them by flex-mer similarity. Seed retrieval stops once seeds from at least four value-blocks have been collected and the seed list exceeds max seed size (default 128). Retrieved seeds are then sorted by taxon identifier, gene identifier and read position.

Seeds are grouped into anchors by taxon and marker gene. Within an anchor, seeds must have consistent spacing in the read and reference, corresponding to ungapped matches between seed positions. Protal extends each anchor along the reference until a mismatch is encountered or another seed is reached. Anchors are ranked by the number of exactly matching positions. For paired-end reads, if the top three anchors are not shared between mates by taxon and gene, protal retrieves additional seeds to recover compatible paired-end anchors.

The top three anchors and any additional recovered anchors are extended into full gapped alignments using WFA2 [16, 17], with mismatch penalty 4, gap-opening penalty 6 and gap-extension penalty 2. Protal then pairs mate alignments and sorts candidate alignment pairs by normalized alignment score. With this penalty scheme, a perfect alignment has score 0, and dividing the score by the mismatch penalty gives a mismatch-equivalent error count. Protal estimates an ANI-like quantity as

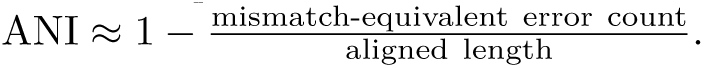

The best-scoring alignment pair is reported.

For each reported alignment, protal stores the number of long-unique and LSU k-mers in the optional SAM columns ZU:i: and ZT:i:, respectively. Protal also reports a mapping-quality score (MAPQ), intended to represent the phred-scaled probability that a read pair is incorrectly mapped. MAPQ decreases when a read pair has multiple candidate alignments with similar scores and increases when one alignment is clearly preferred. Protal follows the minimap2 MAPQ formulation [25], omitting the chaining factor:

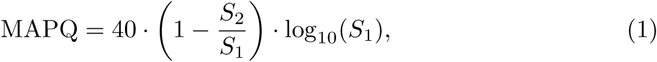

where *S*_1_ and *S*_2_ are the best and second-best alignment bit-scores (computed with match reward +2 and penalties of 3, 1 and 2 for mismatches, gap openings and gap extensions), so that larger values indicate better alignments. When *S*_1_ = *S*_2_, MAPQ is set to 0.

Protal outputs one row of alignment-derived metrics for every species with at least one assigned read (Suppl. Table S1).

### Species detection using machine learning

No single alignment metric reliably distinguishes true species presence from spurious cross-mapping, particularly for low-abundance or closely related taxa. Protal therefore integrates per-species alignment-derived metrics (Supplementary Table S1) in a random-forest classifier [18] that predicts whether a species is present in a metagenome.

Training data were generated from synthetic metagenomes simulated from annotated GTDB r226 source genomes. We used simulate metagenomes (https://github.com/4less/protal) to generate six datasets comprising 581 samples in total. The simulated datasets varied in sequencing depth, number of species, number of samples and abundance distribution. Community composition was further biased towards challenging cases by forcing inclusion of selected taxa, including *Bacteroides ovatus*, and by requiring predefined numbers of species from selected genera, including *Collinsella*, *Escherichia* and *Bacteroides*. For simulations containing multiple strains per species, a strains-per-species parameter such as 0.8,0.8,0.8 was interpreted as sequential probabilities for adding a second, third and additional strains after the first strain, which was always included. Reads assigned to a species were then split among the selected strains.

Protal was run on these synthetic datasets, and the known community composition was used as the gold standard. The classifier was trained on one row per species– sample pair with at least one assigned read. We used a fixed 80/20 train/test split, 256 trees and a maximum of 128 leaf nodes per tree. The number of variables considered at each split (mtry) was tuned using 5-fold cross-validation. The final model was fitted on the training split, and performance was evaluated on both training and test splits. The trained model and feature importances were saved.

Expected gene presence was used as one of the classifier inputs. For a taxon *t*, the expected number of marker genes observed after mapping *reads*(*t*) reads across *genes*(*t*) available marker genes was calculated as:

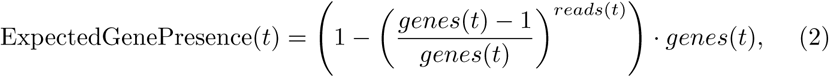

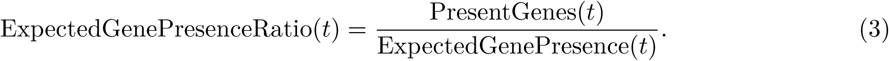

For example, if a taxon has 120 marker genes and 350 reads mapped in total, the expected number of observed genes is 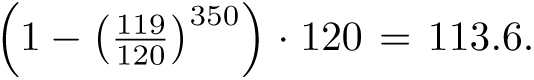. If 50 genes are observed, the expected gene presence ratio is 50*/*113.6 = 0.44.

### Abundance calculation

For each detected species, protal estimates relative abundance from per-gene read depth. Read depth (vertical coverage) for each marker gene is calculated as the total aligned read length divided by marker-gene length, following the Lander–Waterman coverage relationship [51]. The species-level abundance estimate is the median read depth across marker genes assigned to that species.

### Reference-guided multiple sequence alignment construction

For species detected across multiple samples, protal reconstructs within-species phylogenies by building one reference-guided multiple sequence alignment (RG-MSA) of marker-gene consensus sequences per species. This enables strain-level comparison across samples without *de novo* assembly or whole-genome alignment.

### Consensus sequence reconstruction

For each species and marker gene, protal groups all read alignments by sample. Alignments with MAPQ *≤* 4 or soft-clip-corrected alignment length below 50 bases are discarded. From the remaining alignments, protal extracts variants relative to the reference marker gene and records vertical base coverage at each reference position. A minimum coverage of 2 is required to call a base at an MSA position.

For multi-allelic positions, protal chooses the allele with the highest sum of base-quality scores. The selected allele must have coverage of at least 2 and either a summed Phred base-quality score of at least 90 or a mean base-quality score of at least 15. For example, if a position contains bases (A, 20), (A, 35), (A, 40), (T, 25), (T, 40), the consensus base is A because its quality sum is 95, compared with 65 for T. This allele passes the quality filter because it has coverage 3 and summed quality 95.

### Reference-guided MSA construction

*De novo* MSA construction becomes computationally expensive for large numbers of sequences. Protal therefore constructs reference-guided MSAs, which scale linearly with the number of samples for a fixed reference length. Reference-guided MSA approaches have also been used by ViralMSA [19] and VIRULIGN [20] to process large viral sequence collections. In protal, this strategy is appropriate because GTDB marker genes are conserved, and retained reads are expected to align closely to the marker-gene reference.

For each reference marker-gene position, protal applies the following rules. If vertical coverage is at least 2 and a variant passes quality control, protal inserts the consensus variant. Insertions are added to the alignment, and deletions are represented as -. If a variant is observed but fails quality control, protal inserts N. If no variant is observed and vertical coverage is at least 2, protal inserts the reference base. Insertions are handled in two passes: first, protal determines the maximum insertion length across samples at each reference position; second, it pads non-inserting samples with - characters to maintain column alignment.

### MSA quality control and filtering

Per-species MSAs were post-filtered using qcmsa.py. The script operates on protal’s native MSA output, consisting of the raw alignment (`.raw.msa.fna`), the RAxML-style gene partition and a per-(sample, gene) metrics table (`.meta.tsv`). The metrics table reports horizontal coverage (hcov), mean depth over covered positions (mean vcov nonzero) and multi-allelicity rate (MRate2), defined as the fraction of positions carrying more than one allele above the variant-coverage threshold. Filtering proceeds in four stages. Reference sequences absent from the metrics table are exempt from sample-level removal.

### Coverage gating

A (sample, gene) cell was retained if horizontal coverage was at least 0.3 and mean depth was at least 1.0*×*. A gene was retained only if more than three samples passed this cell-level filter. Cells failing coverage within retained genes were gap-filled with - rather than removed. Because removed genes and gap-filled cells were excluded from subsequent multi-allelicity statistics, all adaptive thresholds were calculated on covered data only.

### Multi-allelicity filtering

Potentially contaminated or mixed-strain samples and unreliable genes were identified using an iterative Tukey interquartile-range outlier rule applied to the number of multi-allelic cells per gene and per sample. A cell was considered multi-allelic if MRate2 was greater than 0. Because MRate2 was sparse, the upper fence, *Q*3+*k·IQR*, was calculated using non-zero counts only. By default, *k* = 1.5, and an item was flagged only if its count exceeded the upper fence and it had at least two bad peers. The fence required at least four non-zero values to be applied. In each iteration, outlier genes were flagged first, sample counts were recomputed on surviving genes, and outlier samples were then flagged. Flagged genes and samples were removed, and the procedure was repeated until no further outliers were found or 100 iterations had been reached. Individual (sample, gene) cells with MRate2 above the Tukey fence calculated over non-zero MRate2 values were masked with gaps.

Optional settings allowed the fence to be calculated over the full distribution including zeros, and absolute cutoffs could flag any gene or sample with multi-allelic signal in at least *N* cells.

### Site cleanup

Within surviving columns, all-missing columns and low-parsimony sites were removed, whereas constant (invariant) sites were retained by default because they inform branch-length estimation; constant-site removal can be re-enabled with `--remove-constant.` Low-parsimony sites were defined as positions at which fewer than two samples differed from the majority base.

### Sequence floor

In the integrated protal pipeline this stage is enabled by default: sequences with fewer than msa min hcov (default 1000) valid non-gap, non-N bases after gene and cell removal were removed. Run standalone, qcmsa.py leaves this stage off unless `--reapply-hcov` is set. This filter was applied to the gene-filtered alignment before site cleanup.

The filtered alignment, recomputed partition coordinates and a machine-readable decision log were written for each species. The decision log records each removed gene, sample, site and cell together with its removal reason. Multi-allelicity filtering was controlled by a single preset: strict (*k* = 1.0, min bad=1), default (*k* = 1.5, min bad=2) or sensitive (*k* = 2.0, min bad=3). All thresholds were individually overridable.

### Benchmarking metrics

We evaluated species detection using sensitivity, precision and F1-score, treating each taxon as present or absent relative to the gold standard. Abundance estimation was evaluated using Bray–Curtis dissimilarity, L2 error and Pearson correlation between predicted and expected abundance vectors. Metrics were calculated both on the union and the intersection of predicted and gold-standard taxa where applicable. Custom scripts used for benchmarking are available at https://github.com/4less/benchpro_rust.

For strain-level benchmarking, we used three complementary classes of metrics: MSA error, strain monophyly and tree similarity to gold-standard whole-genome phylogenies. MSA error was calculated by comparing reconstructed marker-gene consensus sequences to the corresponding source genome sequence. For monophyly, samples simulated from the same source genome were expected to form a monophyletic group. For each source genome, we identified the lowest common ancestor (LCA) of all corresponding sample tips and calculated monophyly as the number of tips from the source genome divided by the total number of tips under that LCA. A score of 1 indicates perfect monophyly; values below 1 indicate that the strain was not fully resolved from neighbouring strains.

All strain-level comparisons were restricted to species that every workflow reconstructed and for which protal’s universal markers carried at least 500 variable alignment columns. Of the 46 gold-standard species, StrainPhlAn 4 produced no alignment for one (*Blautia A* sp003471165, which has no counterpart in MetaPhlAn 4’s SGB-to-GTDB mapping), and five further species were removed because protal retained fewer than 500 variable columns for them (*Bacteroides ovatus*, 104; *Lachnospira eligens A*, 257; *Bifidobacterium adolescentis*, 367; *Bacteroides thetaiotaomicron*, 387; *Collinsella* sp003466125, 421). These alignments are full length (40–65 kb) but nearly invariant, so they carry too little phylogenetic signal to score any tool on; the threshold was applied to the species set, so all tools were compared on the same 40 species.

Gold-standard phylogenies were constructed from source genomes using Roary [32] with `--mafft` [33] on Prokka [34] gene predictions. For each species, inferred and reference trees were restricted to shared tips, requiring at least four shared tips. Tree similarity was quantified using branch-score distance [35] and path difference [36]. We additionally calculated Robinson–Foulds distance, weighted Robinson–Foulds distance and Kuhner–Felsenstein distance using the phangorn package [52] for supplementary analyses.

### Datasets and benchmarks

We benchmarked protal across three tasks: read-alignment accuracy, species-level taxonomic profiling and strain-level phylogeny reconstruction. Each task used dedicated datasets and comparison tools as described below.

### Read-alignment benchmark

To evaluate read-alignment accuracy, we simulated reads from all GTDB r214 genomes (*n* = 394,932), including non-representative genomes, using ART Illumina 2.5.8 [22] with parameters: [`-nf 0-p-i reference.fna-o output-l 150-ss HS25-f 5.0-m 200-s 10.`] This yielded 2 *×* 944,933,468 paired-end reads. Because some marker genes are shorter than 200 nt, 128 N characters were added to both ends of each marker gene before simulation to allow read pairs to span the full marker-gene sequence.

We aligned simulated reads against representative GTDB r214 marker genes using protal and four established short-read aligners: Bowtie2 2.5.3 [23], BWA-MEM2 2.2.1 [24], minimap2 2.28-r1209 [25] and Strobealign 0.13.0 [26]. Alignment parameters were --very-sensitive-p 16 for Bowtie2;-t 16 for BWA-MEM2, using bwa-mem2.avx to enforce AVX usage;-ax sr-t 16 --secondary=no for minimap2;-t 16-U for Strobealign; and --max score ani 0.9-u 256-s 128-t 16 for protal. Reference databases for all tools were built from GTDB r214 bacterial species-representative marker genes.

An alignment was counted as a true positive if the read mapped to a marker gene from the representative genome of the species cluster from which the read was simulated. False positives were alignments to any other species cluster. False negatives were calculated relative to the maximum number of aligned reads, correct or incorrect, observed across all aligners. True-positive and false-positive rates were calculated across MAPQ thresholds.

### Computational setup for alignment benchmarks

Alignment speed and memory benchmarks were run on an exclusively reserved compute node with an AMD EPYC 7713P CPU and 512 GB RAM. Each tool was run twice, with the first run used to warm filesystem caches and the second run used for reporting. The input comprised a subset of the simulated marker-gene reads totalling 7.9 GB of uncompressed paired-end read data.

### Species-level profiling benchmarks

We benchmarked species-level profiling using CAMI synthetic metagenomes, a rare-species benchmark and a high-complexity benchmark.

### CAMI synthetic metagenomes

We used the “Toy Human Microbiome Project Dataset” (CAMI Human), “Toy Mouse Gut Dataset” (CAMI Mouse) and “Marine” dataset from the second CAMI challenge [27, 28], available at https://data.cami-challenge.org/. CAMI Human comprises five body-site datasets: Airways, Gastrointestinal, Oral, Skin and Urogenital. Gold-standard profiles were converted to GTDB r226 by running GTDB-Tk v2.3.2 [29] on the source genomes and transferring abundances from CAMI gold-standard files. For CAMI Human, abundances were taken from abundance.tsv; for CAMI Mouse, from distributions/distribution.txt; and genome identifiers were mapped using genome to id.tsv.

### Rare-species benchmark

We generated a rare-species dataset (R.Sp.) to test detection of species represented by only one genome in GTDB r214. From the HumGut database [30], we selected 958 isolate genomes from 364 species with only one representative genome in GTDB r214. Selected genomes were subsequently re-annotated with GTDB-Tk against GTDB r226 for evaluation. We simulated 20 metagenomes, each containing 200 species represented by one randomly selected genome. Vertical genome coverage values were sampled from a distribution bounded between 0.05 and 70.

Paired-end reads of length 2 *×* 150 bp were simulated using ART Illumina [22] with parameters: [`-p-l 150-ss HS25-m 200-s 10.`] This dataset was used only for species-level profiling.

### High-complexity benchmark

We additionally generated a high-complexity synthetic metagenome benchmark containing approximately 10,000 species per sample to evaluate profiler behaviour under extreme community complexity.

### Taxonomic profiling tools

We compared protal v0.6.0, run in both its full and reduced (“protal-mini”) database configurations, with MetaPhlAn 4 v4.2.4 [6], mOTUs v4.0.4 [7], Kraken2 v2.17.1 with Bracken v3.1 [8, 31], and sylph v0.9.0 [9]. MetaPhlAn 4 was run with database mpa vJan21 CHOCOPhlAnSGB 202103 and default parameters. MetaPhlAn 4 uses Bowtie2 [23] with `--sensitive` to align reads against species-specific markers. mOTUs was run using its recommended settings. Kraken2 was run against a custom database built from all GTDB r226 species-representative genomes, producing native GTDB r226 output. Bracken abundance estimation was run with read threshold-t 450. Sylph was run using sylph profile with default parameters against two GTDB r226 databases built at subsampling rates *c* = 200 and *c* = 1000 (`gtdb-r226-c200` and `gtdb-r226-c1000`), which trade sensitivity against speed.

Profilers differ in how they report clades they cannot name: MetaPhlAn 4 aggregates them into a single UNCLASSIFIED entry, whereas mOTUs4 emits one place-holder species per unnamed marker-gene cluster (`s__Unknown <genus> mOTUv4.0 *`), resolved to genus but without a species name. Because such an entry can never match a gold-standard species, scoring it as a species prediction would penalise a profiler for being explicit about what it could not resolve. We therefore excluded predictions that are unnamed at the rank being evaluated from the true-positive/false-positive accounting at that rank, for all tools alike. These predictions still count at ranks where they carry a name, so results at genus level and above are identical either way. In our benchmark this affected only mOTUs4, for which 3,007 of 6,174 species-level false positives were such placeholders; 99.9% of them fell within a genus that mOTUs4 had correctly detected in the same sample.

Sylph and Kraken2+Bracken natively report profiles in GTDB taxonomy, as both were run against GTDB r226 databases. For the remaining tools, outputs were converted to GTDB taxonomy where required. MetaPhlAn 4 profiles were mapped to GTDB using sgb to gtdb profile.py; NCBI-formatted output was generated using --CAMI format output. mOTUs output was mapped to GTDB using the corresponding mOTUs-to-GTDB mapping file and a custom script.

### Strain-level phylogeny benchmark

We generated the STRAIN46 dataset to benchmark strain-level phylogeny reconstruction in a setting resembling longitudinal or multi-sample strain tracking, where some samples share the same source strain. Source genomes were selected from HumGut [30]. We filtered HumGut for isolate genomes and selected 50 species with 15–60 isolate genomes each. Because the original HumGut taxonomy used GTDB r95, we re-annotated the pool of 1,428 isolate genomes with GTDB r214. After re-annotation, these genomes spanned 67 GTDB r214 species. Species with fewer than 15 genomes were removed, leaving 1,346 genomes across 46 species.

We simulated 200 metagenomes. Each sample contained one strain from each of the 46 species. For each sample and species, one genome was selected randomly, and vertical abundance was drawn from a negative-binomial distribution scaled between 1 and 50. Paired-end reads (2 *×* 150 bp) were simulated with ART Illumina using: [`-ss HS25-i <input.fna>-p-l 150-f-m 200-s 10-o <output_prefix>.`]

For each species, we constructed a gold-standard core-genome MSA using Roary [32] with `--mafft` [33] on Prokka [34] gene predictions from the source genomes. Gold-standard phylogenies were inferred using IQ-TREE (v3.1.1) [21] with: [`iqtree-s-fast-m GTR.`]

### Strain-level profiling

We compared protal’s strain-level output with StrainPhlAn 4 v4.2.4 [6]. For StrainPhlAn 4, marker regions were first extracted from each sample using sample2markers.py; database marker regions were extracted from all SGBs using extract markers.py. Per-species MSAs were then generated using StrainPhlAn 4 with `--marker in n samples 20.` Protal was run with default parameters to generate taxonomic profiles and per-species marker-gene MSAs. Phylogenetic trees were inferred from protal MSAs using IQ-TREE (v3.1.1) [21] with: [iqtree-s-fast-m GTR.]

### Strain-level runtime benchmark

For strain-level runtime benchmarking, we ran protal and MetaPhlAn 4+Strain-PhlAn 4 on increasing subsets of STRAIN46, using the first *n* samples for *n* = 4, 8, 16, 32, 64 and 128. Protal, MetaPhlAn 4 and the StrainPhlAn 4 commands `extract markers.py and sample2markers.py` were run sequentially with 32 cores and no other programs running concurrently. Peak memory was recorded as the maximum resident set size across all stages.

StrainPhlAn 4 was run with: [`--breadth thres 50 --marker in n samples 50 --sample with n markers 5.`]

## Notes

### Competing Interest Statement

The authors have declared no competing interest.

### Summary of Updates

Species benchmarking updated to better handle profilers with results unresolved to species level (Fig. 2). Strain MSA errors updated based on additional filtering (Fig 3b) and mean monophyly updated (Fig 3d). Revisions in text on these same topics also. Supplementary Tables added.

